# CTDP: Identifying cell types associated with disease phenotypes using scRNA-seq data

**DOI:** 10.1101/2025.10.16.682537

**Authors:** Chonghui Liu, Murong Zhou, Zhidong Wang, Chen Yang, Yan Zhang, Zhongjun Jiang, Yu-Hang Yin, Zeyu Luo, Tianjiao Zhang, Guohua Wang, Lei Yuan

## Abstract

Single-cell RNA sequencing enables transcriptome-wide analysis at single-cell resolution, offering unprecedented insights into cellular heterogeneity across biological conditions. However, accurately comparing transcriptomic distributions of cells from distinct biological states, such as healthy versus diseased individuals, remains challenging. To address this, we developed CTDP, a robust and interpretable computational framework that identifies disease phenotype-associated cell types of interest by integrating Lasso-regularized logistic regression with permutation testing. Through comprehensive evaluations on both simulated and real-world datasets, including melanoma immunotherapy, COVID-19 severity, and liver cirrhosis, CTDP consistently outperformed existing methods such as DA-seq, scDist, and PENCIL in both accuracy and robustness. In melanoma, CTDP uncovered immune-responsive clusters and revealed transcriptional regulators like PTPRC, CREM, and JUNB linked to immunotherapy efficacy. In COVID-19, it identified critical severity-associated cell types, such as B cells, NK cells, epithelial cells, and macrophages, which contribute to dysregulated immune responses and inflammation in severe cases. These results highlight CTDP’s power in uncovering disease-relevant cell populations and its potential to advance precision medicine through single-cell analysis.

## Introduction

Single-cell RNA sequencing (scRNA-seq) has revolutionized our understanding of cellular heterogeneity by enabling transcriptome-wide measurements at single-cell resolution across a wide range of biological systems^1^. This powerful technology provides a unique opportunity to investigate how gene expression varies across individual cells under diverse conditions, such as healthy versus diseased states, treatment versus control, or responders versus non-responders to therapy^2, 3^. Despite its transformative potential, extracting meaningful phenotype-related insights from scRNA-seq data remains a major challenge, primarily due to its high dimensionality, sparsity, and technical noise. Therefore, there is a critical need for robust computational methods capable of accurately identifying phenotype-associated cell populations in such complex datasets.

A widely adopted approach for detecting such populations involves clustering all cells across conditions in an unsupervised manner^4^. Once clusters are defined, the relative proportions of cells from each condition within each cluster are compared to their global proportions in the dataset. Clusters that exhibit statistically significant deviations are regarded as differentially abundant between conditions. This general framework has proven effective across a range of biological applications, including characterizing immune responses across disease severities following viral infection^5^, comparing therapeutic outcomes in cancer immunotherapy^6^, and studying cell remodeling in chronic inflammatory conditions such as inflammatory bowel disease^7^.

In addition to this general strategy, several computational methods have been developed, including DA-seq^4^, scDist^8^, and PENCIL^9^. These tools aim to identify differentially abundant or differentially expressed cell populations between phenotypic groups. While these methods have facilitated important discoveries, they often exhibit limitations. For example, some struggle to detect weak phenotype signals, others offer limited support for detecting multiple associated clusters, and many lack rigorous statistical controls for false discoveries. These limitations highlight the ongoing need for computational frameworks that are not only accurate and sensitive but also interpretable and statistically robust.

To address this need, we developed CTDP (Identification of <u>C</u>ell <u>T</u>ypes associated with <u>D</u>isease <u>P</u>henotypes), a novel framework based on Lasso-regularized logistic regression, augmented with permutation testing and false discovery rate correction. CTDP is designed to robustly identify phenotype-associated cell clusters from high-dimensional single-cell data. Through extensive evaluations on both synthetic and real-world datasets, CTDP consistently outperformed existing methods in terms of accuracy and robustness. Notably, it effectively uncovered disease-relevant cell types and regulatory mechanisms in studies of melanoma immunotherapy, COVID-19, and liver cirrhosis. These findings underscore CTDP’s broad applicability and its potential to drive advances in precision medicine by providing deeper insights into cell-type-specific contributions to disease.

## Results

### Overview of CTDP

To systematically identify cell clusters associated with disease phenotypes from single-cell transcriptomic data, we developed CTDP (Identification of <u>C</u>ell <u>T</u>ypes associated with <u>D</u>isease <u>P</u>henotypes)—a computational framework that integrates Lasso-regularized logistic regression with rigorous statistical inference. As illustrated in Fig. 1, CTDP operates on two primary inputs: (i) scRNA-seq data annotated with cluster identities, and (ii) sample-level disease phenotype labels, which are propagated to individual cells (Fig.1, top). The workflow begins with data preprocessing and clustering using Seurat, followed by one-hot encoding of cluster assignments into a binary matrix and phenotype labels into a binary response vector. A Lasso-regularized logistic regression model is then fitted to these encoded data, leveraging the sparsity-inducing property of Lasso to identify a minimal set of cell clusters predictive of the phenotype, with the optimal regularization parameter (λ) determined via cross-validation to ensure model generalizability (Fig.1, bottom).

**Fig. 1.**
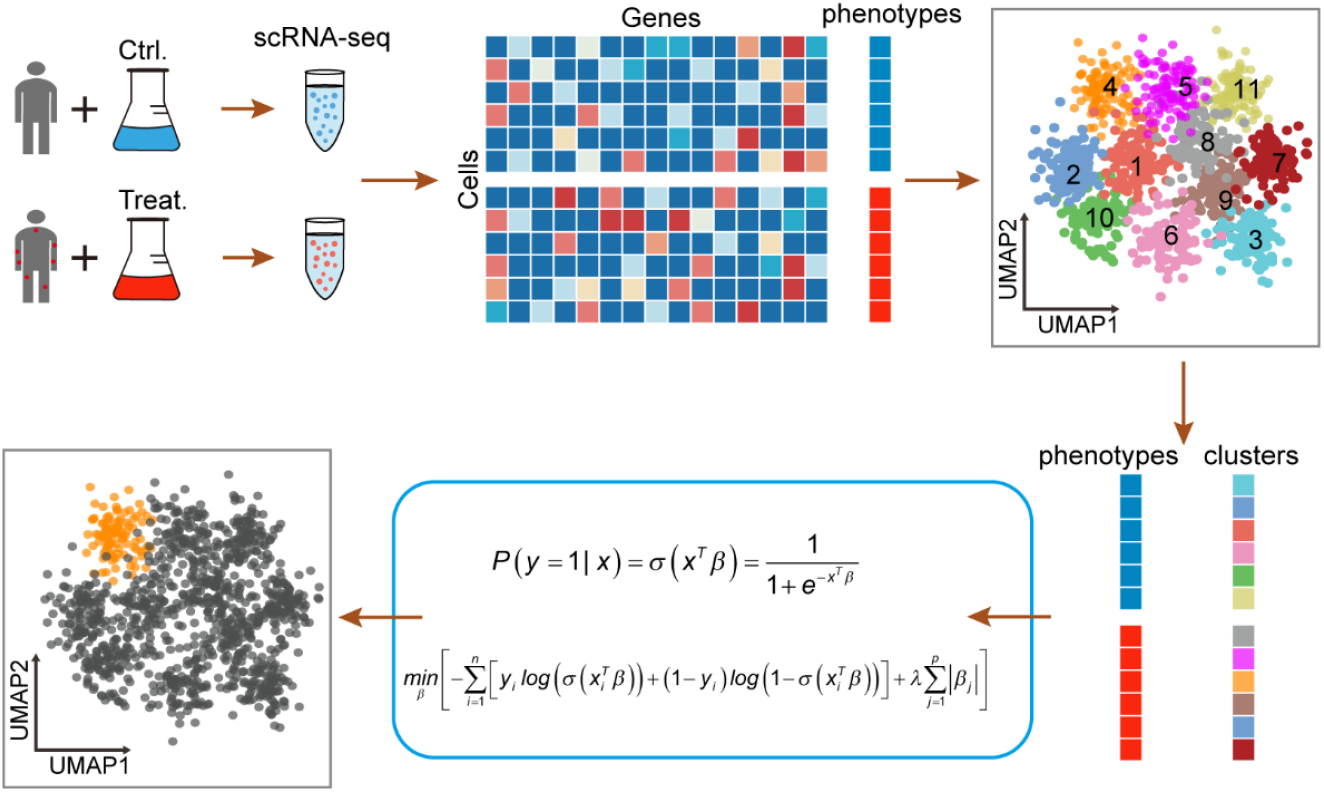
The workflow of the CTDP. CTDP is a computational framework designed to identify cell clusters associated with disease phenotypes from scRNA-seq data. Input data include: (i) single-cell transcriptomic profiles, and sample-level disease phenotype labels, where each cell is annotated with the phenotype of the sample it originated from (top left and top center). The CTDP workflow consists of three major steps. (1) Clustering and data encoding: The input scRNA-seq data are first analyzed using Seurat to perform standard preprocessing and clustering (top right). The resulting cluster assignments are then one-hot encoded into a binary matrix, and phenotype labels are encoded as a binary response vector (bottom right). (2) Model fitting: A Lasso-regularized logistic regression model is used to assess the association between each cluster and the disease phenotype, with the regularization parameter selected via cross-validation (bottom center). (3) Statistical inference: Permutation testing is conducted to compute empirical p-values for each cluster, and the Benjamini–Hochberg procedure is applied to control the false discovery rate (bottom left).

To evaluate the statistical significance of each cluster selected by the model, CTDP performs permutation testing by randomly shuffling phenotype labels and recalculating regression coefficients under the null distribution. Empirical p-values are then derived and adjusted for multiple comparisons using the Benjamini–Hochberg procedure to estimate false discovery rates (FDR). This integrative framework enables accurate and interpretable identification of phenotype-associated cell clusters while effectively controlling false positives in high-dimensional single-cell datasets.

### Performance of CTDP on simulated data with a single disease-associated cell cluster

To comprehensively evaluate the effectiveness of CTDP in identifying phenotype-associated cell clusters, we designed a series of simulation experiments and conducted comparative analyses with three existing methods: DA-seq, scDist, and PENCIL. We based our simulations on a real human immune cell scRNA-seq dataset comprising 4,669 cells^10^. To simulate disease-related phenotypic heterogeneity, we generated multiple synthetic datasets by assigning phenotype labels according to predefined rules.

First, standard preprocessing steps were applied to the scRNA-seq dataset, including the removal of low-quality cells, normalization of gene expression values, and cell clustering. Among the resulting clusters, we selected one specific cluster (Cluster 1) as the true phenotype-associated cell cluster (Fig. 2A). We then generated phenotype labels with a mixing ratio of 0.1: 90% of cells in Cluster 1 were assigned as “disease” and 10% as “control,” while cells in other clusters were assigned phenotype labels uniformly at random (Fig. 2B). This configuration reflects a realistic scenario where one subpopulation is enriched in the disease condition.

**Fig. 2.**
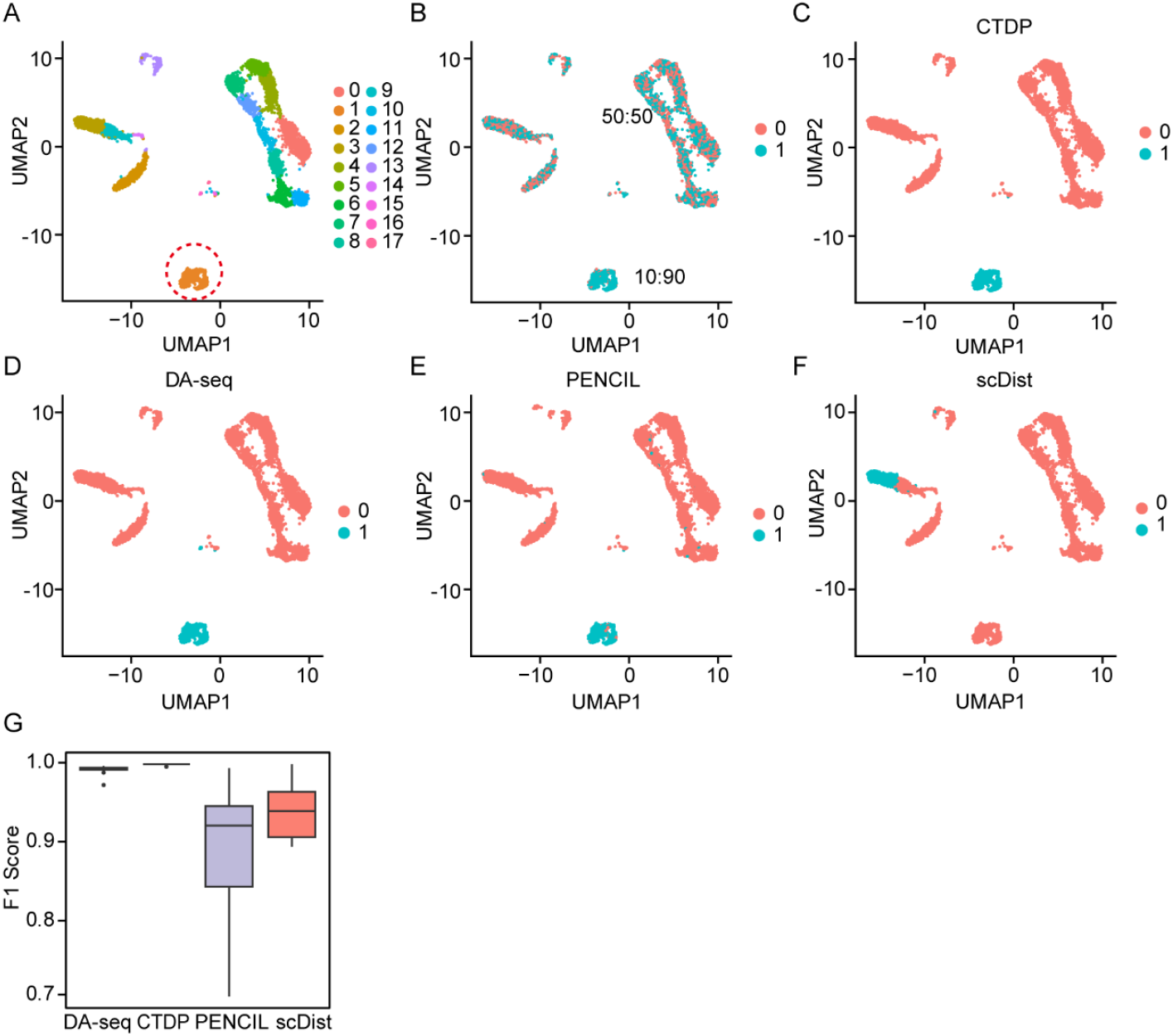
Evaluation of CTDP’s prediction in simulations. (A) UMAP plot of cell type distribution in the simulated data, where the cell cluster highlighted with a red circle is defined as the phenotype-associated cluster of interest, and other clusters serve as controls. (B) The phenotype-associated cluster of interest is assigned two phenotype labels based on a mixing ratio of 0.1, while the remaining cells are assigned phenotype labels evenly, as shown in the proportions. (C–F) Predictions of phenotype-associated cell types by four methods: CTDP (C), DA-seq (D), PENCIL (E), and scDist (F). (G) Boxplot comparing the performance of the four methods on simulated data, with one phenotype-associated cluster randomly selected in each of 10 trials (n = 10). In the boxplots, the center line represents the median, and the box boundaries indicate the upper and lower quartiles.

Using the resulting synthetic dataset, we provided the single-cell expression matrix and the simulated phenotype labels as input to CTDP, DA-seq, scDist, and PENCIL, and assessed their performance in identifying the phenotype-associated cluster. To quantitatively compare the methods, we adopted the F1 score as the main evaluation metric. This metric balances both the accuracy and completeness of detection, providing a comprehensive measure of a method’s performance in identifying disease-relevant cell types.

As shown in Fig. 2C-F, CTDP achieved superior performance in detecting the true phenotype-associated cell cluster. To assess the robustness of the methods, we repeated the simulations ten times, each time selecting a different cluster as the disease-relevant target. In all repetitions, CTDP consistently outperformed DA-seq, scDist, and PENCIL (Fig. 2G). DA-seq showed competitive but slightly inferior performance, while PENCIL and scDist demonstrated significantly lower F1 scores. These results highlight the superior sensitivity and specificity of CTDP in identifying phenotype-associated cell clusters, even in challenging scenarios with imbalanced label distributions.

### Performance of CTDP on simulated data with multiple disease-associated cell clusters

To further evaluate the flexibility and robustness of CTDP in complex biological scenarios, we extended the simulation framework to include multiple phenotype-associated cell clusters. Using the same scRNA-seq dataset consisting of 4,669 human immune cells, we randomly selected three distinct clusters (cluster 2, cluster 5, and cluster 13) as the true phenotype-associated cell clusters (Fig. 3A, B). This setting aims to mimic more realistic disease conditions, where multiple cell subpopulations may collectively contribute to the disease phenotype.

**Fig. 3.**
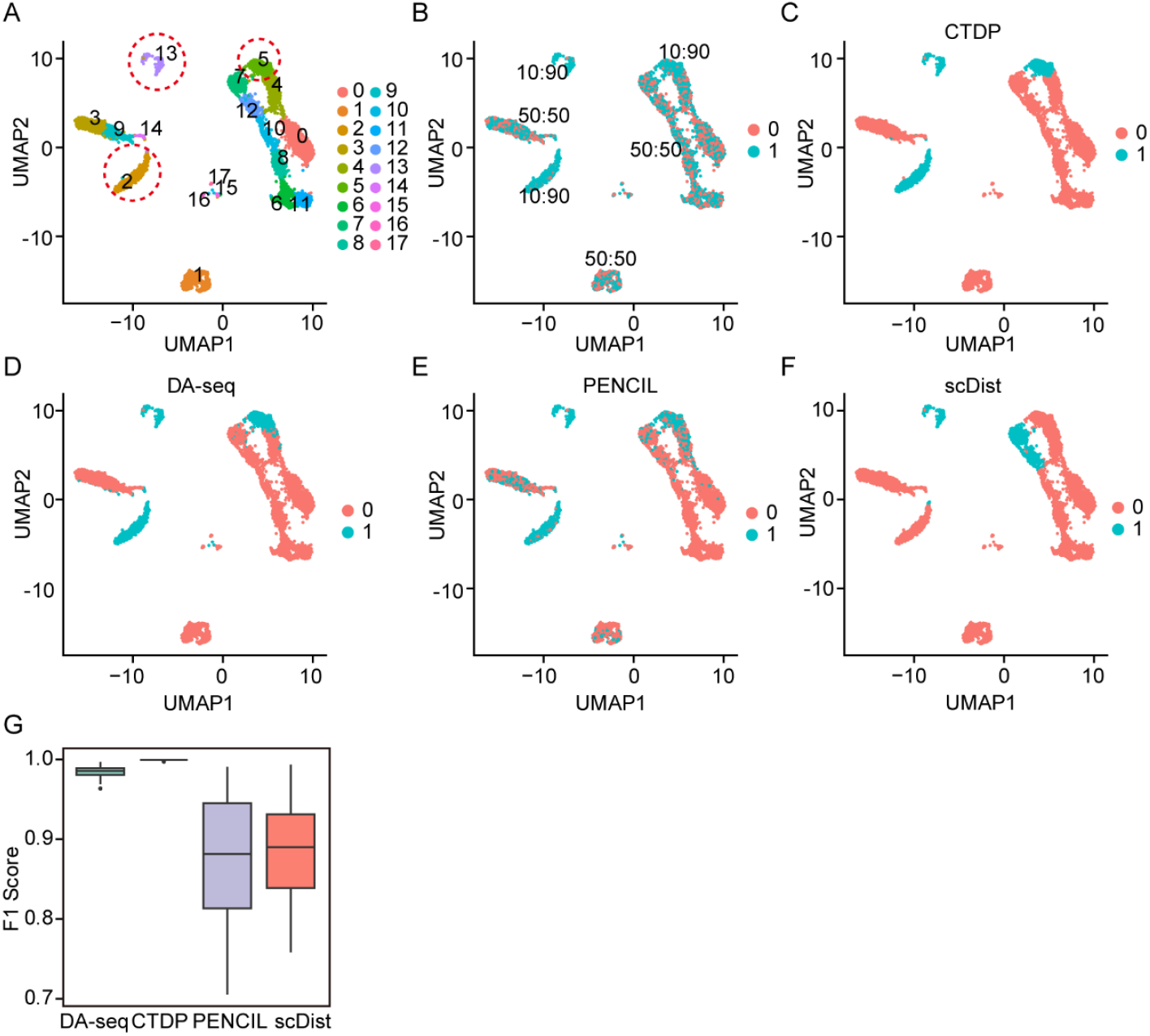
Evaluation of CTDP’s prediction performance on simulated datasets with multiple phenotype-associated cell clusters of interest. (A) UMAP plot of cell type distribution in the simulated data, where the cell cluster highlighted with a red circle is defined as the phenotype-associated cluster of interest, and other clusters serve as controls. (B) The phenotype-associated cluster of interest is assigned two phenotype labels based on a mixing ratio of 0.1, while the remaining cells are assigned phenotype labels evenly, as shown in the proportions. (C–F) Predictions of phenotype-associated cell types by four methods: CTDP (C), DA-seq (D), PENCIL (E), and scDist (F). (G) Boxplot comparing the performance of the four methods on simulated data, where three phenotype-associated clusters are randomly selected in each of 10 trials (n = 10). In the boxplots, the center line indicates the median, and the box edges represent the upper and lower quartiles.

As shown in Fig. 3C-F, CTDP achieved the best predictive performance among all methods, followed by DA-seq, whereas scDist showed the lowest accuracy. These findings demonstrate the superior ability of CTDP to detect multiple disease-relevant subpopulations simultaneously. To assess the robustness of the results, we repeated the simulation 10 times, each time randomly selecting three different clusters as phenotype-associated subpopulations. Consistently, CTDP outperformed the other methods across all runs (Fig. 3G), confirming its strong adaptability and predictive accuracy under multi-cluster settings.

### Discovery of immunotherapy-responsive cell clusters by CTDP

To demonstrate the applicability of CTDP in real-world clinical scenarios, we applied the method to a scRNA-seq dataset containing 16,291 immune cells derived from tumor samples of 48 melanoma patients treated with immune checkpoint inhibitors (ICIs) (Fig. 4A, B). The dataset was designed to investigate the immune cell landscape associated with therapeutic response. Based on the clinical response to ICIs, CTDP successfully identified seven cell clusters (clusters 0, 2, 4, 5, 7, 14, and 15) that were significantly enriched in responders (Fig. 4C). The identification of these immune-responsive subpopulations provides valuable insights into the cellular basis of ICI efficacy.

**Fig. 4.**
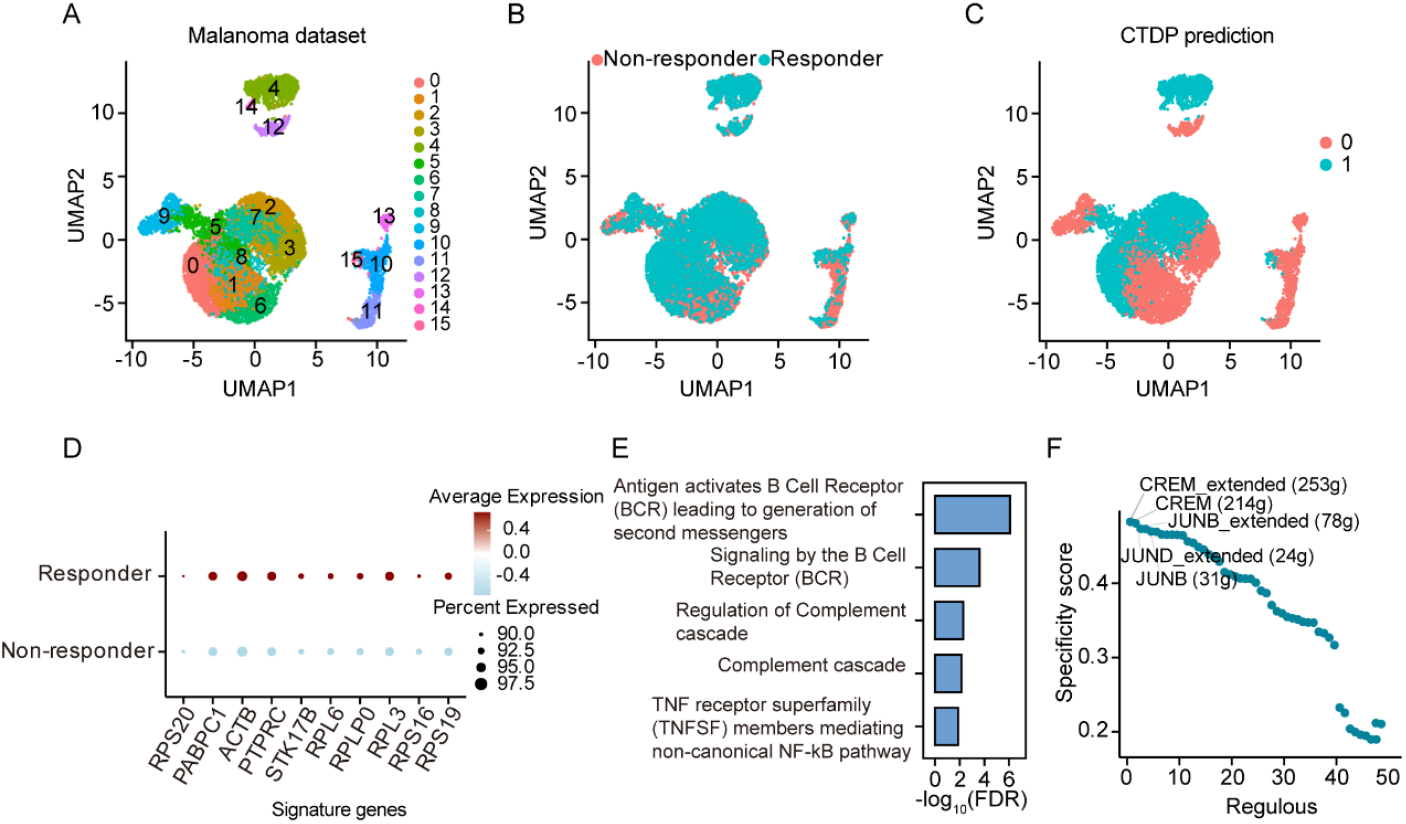
CTDP identifies cell clusters associated with immune responders. (A) UMAP plot showing the cellular composition of the melanoma scRNA-seq dataset, with cells colored by cluster labels. (B) Cells colored by immunotherapy response status. (C) UMAP plot displaying CTDP prediction results, where 1 represents the predicted immune-responsive cluster and 0 represents the others. (D) Dot plot showing the top 10 differentially expressed genes between cells in the phenotype-associated cluster related to immunotherapy responders and other cells, with color intensity indicating the average gene expression level. (E) Top five Reactome pathways enriched in highly expressed genes within CTDP-identified cells. (F) Ranking of regulators based on RSS in the CTDP-identified cells associated with immunotherapy responders.

To further explore the transcriptional programs underlying immunotherapy response, we performed differential gene expression analysis between the immune-responsive clusters identified by CTDP and the remaining non-responder clusters (Supplementary Table 1). The top 10 differentially expressed genes are shown in Fig. 4D. Notably, several genes exhibited expression patterns closely associated with immune responsiveness. In particular, PTPRC (encoding CD45) was significantly upregulated in responder-enriched clusters. Previous studies have reported that PTPRC expression is positively correlated with the expression of immune checkpoint molecules and that PTPRC knockdown promotes melanoma cell migration, invasion, and proliferation^11^. These findings suggest that PTPRC plays a critical regulatory role in the response to ICIs and may serve as a promising biomarker for treatment prediction.

In addition to individual gene markers, we investigated the functional pathways enriched in the immune-responsive clusters. Gene set enrichment analysis revealed that highly expressed genes in these clusters were significantly involved in key immune-related pathways, including “antigen activates B cell receptor leading to generation of second messengers”, “signaling by the B cell receptor”, and “regulation of the complement cascade”, among others (Fig. 4E and Supplementary Table 2). These pathways have been shown to regulate immune cell infiltration, which is essential for enhancing the efficacy of checkpoint-based immunotherapy^12, 13^. The identification of these functionally relevant pathways further supports the mechanistic importance of the clusters revealed by CTDP in shaping the therapeutic response to ICIs.

Moreover, transcription factor (TF) enrichment analysis indicated that the immune-responsive clusters were regulated by specific TFs, particularly CREM and JUNB (Fig. 4F and Supplementary Table 3). Both TFs are essential regulators of immune cell activation, proliferation, and functional modulation. CREM and JUNB have been implicated in the orchestration of T cell responses and immune homeostasis, and may directly influence the efficacy of immune checkpoint blockade therapy^14, 15^. These TFs represent potential regulatory nodes governing immune responsiveness in the tumor microenvironment.

In summary, CTDP not only accurately identified cell subpopulations associated with response to immunotherapy, but also uncovered key transcriptional signatures and regulatory factors underlying treatment efficacy. The highlighted genes and TFs, including PTPRC, CREM, and JUNB, may serve as novel biomarkers and therapeutic targets for personalized immunotherapy. These results illustrate the power of CTDP in dissecting the immune mechanisms of treatment response and offer valuable insights for the development of precision immunotherapeutic strategies.

### Cell types associated with COVID-19 severity revealed by CTDP

To identify cell clusters associated with COVID-19 severity, we applied the CTDP method to single-cell RNA sequencing data from nasopharyngeal swabs, protected specimen brushes, and bronchoalveolar lavage fluid samples collected from 19 COVID-19 patients, including 8 with moderate and 11 with critical disease severity (Fig. 5A, B). To ensure consistency in cell-type distribution across conditions, we performed cell-type-level downsampling on moderate cases. CTDP analysis revealed 13 cell types specifically enriched in patients with critical illness, including B cells, ciliated cells, ionocytes, interferon-γ-responsive cells, mast cells, neutrophils, natural killer cells, macrophages, epithelial cells, dendritic cells, squamous cells, and regulatory T cells (Fig. 5C). These cell populations are key participants in antiviral defense, immune regulation, and inflammation^16-18^. In severe cases, however, their functions may become dysregulated, leading to immune exhaustion, immune evasion, or cytokine storms, all of which are hallmarks of critical COVID-19 pathophysiology^19-21^.

**Fig. 5.**
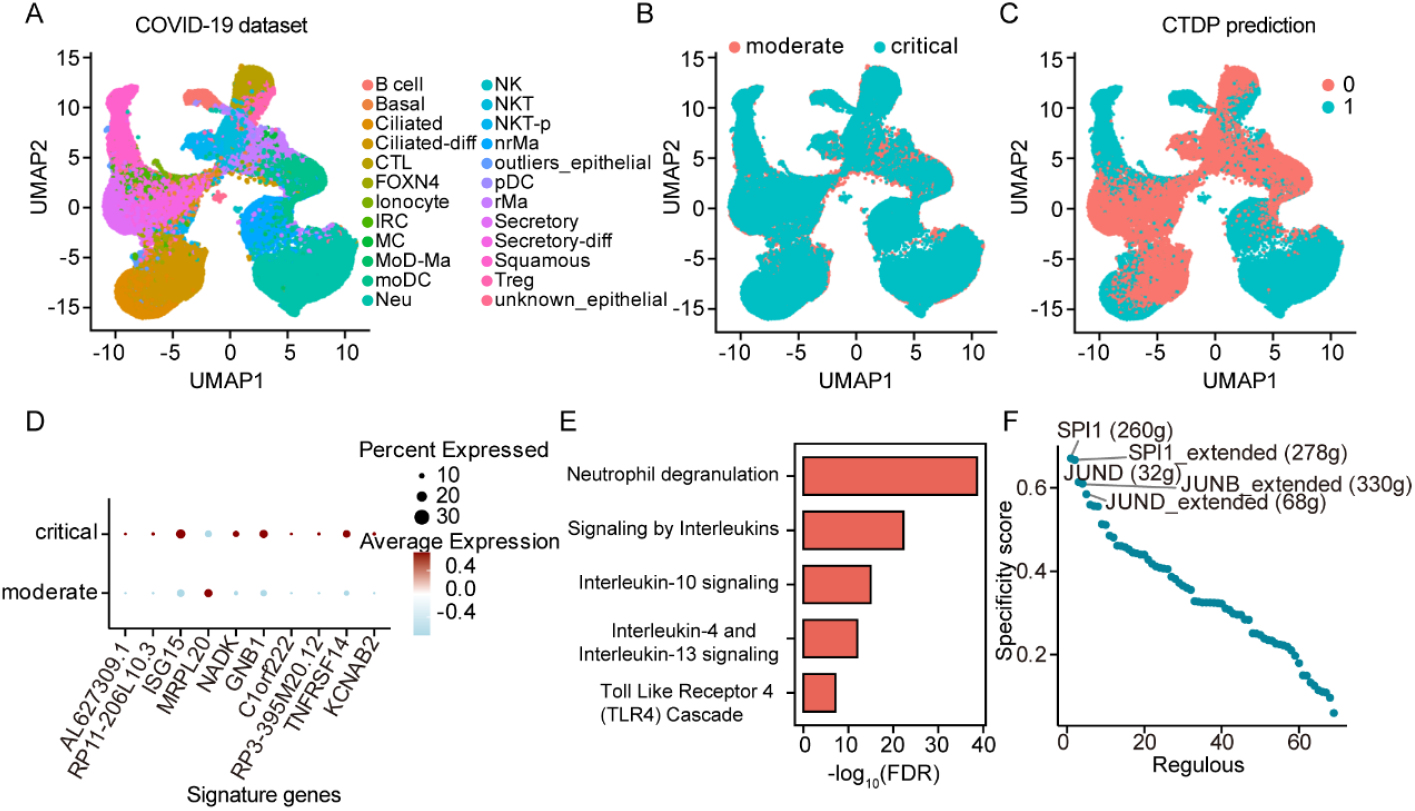
CTDP identifies cell types associated with severe COVID-19 phenotypes. (A) UMAP plot showing the cellular composition of the scRNA-seq dataset from COVID-19 patients, with cells colored by cell type labels. (B) Cells colored by disease status. (C) UMAP plot displaying the CTDP prediction results, where 1 represents the predicted phenotype-associated cell types and 0 represents controls. (D) Dot plot showing the top 10 differentially expressed genes between cells of phenotype-associated cell types related to severe COVID-19 and other cells, with dot color intensity representing the average gene expression level. (E) Top five Reactome pathways enriched in highly expressed genes in cells identified by CTDP. (F) RSS-based ranking of transcriptional regulators in CTDP-identified cells associated with the severe COVID-19 phenotype.

To further investigate molecular features of the critical COVID-19-associated cell populations, we performed differential expression analysis between these cell subsets and the remaining cells (Supplementary Table 4). We identified several highly upregulated genes, including ISG15, NADK, and GNB1, which are involved in antiviral immune responses and metabolic regulation (Fig. 5D)^22-24^. These transcriptional signatures provide insights into the disturbed immune response and altered metabolic landscape in critically ill COVID-19 patients.

Gene set enrichment analysis of the highly expressed genes revealed significant involvement in immune and inflammatory pathways, such as neutrophil degranulation, signaling by interleukin, interleukin-4 and interleukin-13 signaling, and toll-like receptor 4 cascade (Fig. 5E and Supplementary Table 5). Dysregulation of these pathways may underlie the excessive immune responses, immune dysfunction, and cytokine storms observed in critical COVID-19, highlighting them as potential therapeutic targets ^25, 26^.

Moreover, we predicted transcription factors specifically regulating the critical COVID-19-associated cell types (Supplementary Table 6). Key regulators included SPI1, JUND, and others (Fig. 5F). For instance, SPI1 (also known as PU.1) is a master regulator of myeloid and lymphoid cell differentiation and has been implicated in macrophage activation^27, 28^. The enrichment of these regulators in the COVID-19-relevant cell populations suggests their potential roles in orchestrating the immune dysregulation observed in severe cases. These findings highlight possible upstream drivers of transcriptional programs underlying disease-associated cellular states, offering mechanistic insights into the host response to SARS-CoV-2 infection.

In summary, CTDP enabled the identification of critical COVID-19-associated cell populations and their regulatory programs, offering mechanistic insights into immune dysfunction and revealing potential molecular targets for therapeutic intervention in severe COVID-19.

### Application of CTDP to uncover cell types involved in liver cirrhosis

We applied the CTDP method to single-cell transcriptomic data derived from non-parenchymal cells in human liver tissues from healthy donors and patients with liver cirrhosis. To eliminate sampling bias, we performed cell-type-level downsampling on healthy donor-derived cells, ensuring that the total number of cells matched between healthy and cirrhotic samples (Fig. 6A, B). CTDP analysis identified several cell types specifically associated with cirrhotic liver tissues, including plasmacytoid dendritic cells, B cells, plasma cells, endothelial cells, mesothelial cells, hepatocytes, and cholangiocytes (Fig. 6C). These cell populations are critically involved in the pathogenesis and progression of liver cirrhosis, participating in immune regulation, inflammation, fibrogenesis, and structural remodeling of the liver^29, 30^. These findings highlight the complex, multicellular nature of cirrhosis progression, which is not driven solely by the dysfunction of individual cell types but also by their dynamic interactions.

**Fig. 6.**
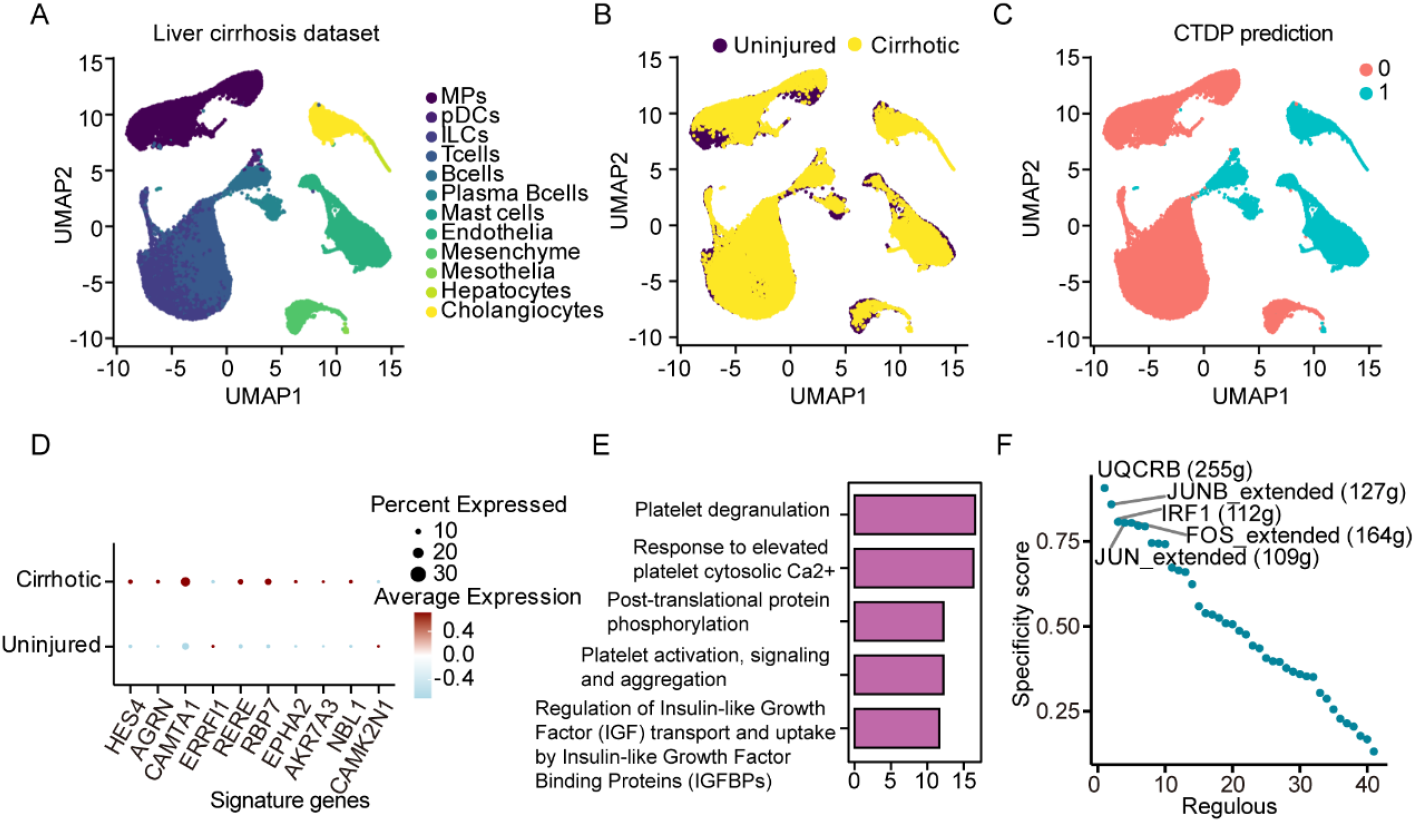
CTDP identifies cell types associated with liver cirrhosis. (A) UMAP plot showing the cellular composition of the scRNA-seq dataset from cirrhosis patients, with cells colored by cell type labels. (B) Cells colored by disease status. (C) UMAP plot displaying the CTDP prediction results, where 1 represents the predicted phenotype-associated cell types and 0 represents controls. (D) Dot plot showing the top 10 differentially expressed genes between cells of phenotype-associated cell types related to cirrhosis and other cells, with dot color intensity representing the average gene expression level. (E) Top five Reactome pathways enriched in highly expressed genes in cells identified by CTDP. (F) RSS-based ranking of transcriptional regulators in CTDP-identified cells associated with cirrhosis.

To investigate the molecular characteristics of the cell subsets identified by CTDP, we analyzed the differentially expressed genes between these subsets and all other cells (Supplementary Table 7). These genes are involved in diverse biological processes relevant to fibrosis and liver pathology. For instance, AGRN and NBL1 are implicated in fibrogenesis, AGRN and EPHA2 in extracellular matrix remodeling, ERRFI1 and CAMTA1 in immune and inflammatory responses, RBP7 and AKR7A3 in metabolic stress and detoxification, and CAMK2N1 and RERE in signaling regulation (Fig. 6D)^31, 32^. These molecular signatures offer valuable insights into the pathophysiology of liver cirrhosis at the single-cell level.

Pathway enrichment analysis revealed that the upregulated genes in cirrhosis-associated cell subsets were significantly enriched in pathways such as platelet degranulation, response to elevated platelet cytosolic Ca^2+^, post-translational protein phosphorylation, platelet activation, signaling and aggregation, and regulation of insulin-like growth factor transport and uptake by insulin-like growth factor binding proteins (Fig. 6E and Supplementary Table 8). These pathways reflect platelet dysfunction, abnormal signal transduction, and disrupted growth factor signaling in liver cirrhosis. Notably, accumulating evidence has shown that platelet responsiveness decreases with increasing severity of liver cirrhosis, although the precise underlying mechanisms remain to be fully elucidated^33, 34^. Moreover, dysregulated protein phosphorylation and insulin-like growth factor signaling may further contribute to cellular dysfunction and tissue injury, potentially representing key drivers of cirrhosis progression and candidate targets for antifibrotic therapy^35, 36^.

Finally, we identified transcription factors that are specifically enriched in cirrhosis-associated cell subsets, including UQCRB, JUNB, FOS, and others (Fig. 6F and Supplementary Table 9). UQCRB, a key component of mitochondrial complex III, is involved in regulating oxidative phosphorylation and reactive oxygen species production, processes known to contribute to oxidative stress and tissue damage during liver fibrosis^37, 38^. FOS, a member of the AP-1 transcription factor family, plays crucial roles in modulating inflammatory responses and fibrogenesis by regulating genes associated with immune activation and cellular proliferation ^39^. The enrichment of these factors in cirrhosis-related cell populations suggests their involvement in driving pathological transcriptional programs underlying liver fibrosis and cirrhosis progression. These findings provide insights into potential molecular mechanisms and therapeutic targets for the treatment of liver cirrhosis.

## Discussion

Understanding how infections, treatments, or biological conditions influence specific cell types is fundamental to elucidating disease mechanisms and informing therapeutic strategies. In this study, we present CTDP, a novel computational framework designed to robustly identify disease phenotype-associated cell clusters from high-dimensional single-cell transcriptomic data. By integrating Lasso-regularized logistic regression with permutation-based hypothesis testing, CTDP effectively balances model sparsity and statistical rigor, offering improved accuracy and interpretability in the detection of disease-relevant cellular populations.

Our simulation studies, which incorporated both single and multiple disease-associated clusters, demonstrate the superior performance of CTDP compared to existing methods such as DA-seq, scDist, and PENCIL. Notably, CTDP achieved consistently higher F1 scores under various conditions, including scenarios with imbalanced label distributions and multiple contributing subpopulations. These results underscore the method’s robustness and generalizability, particularly in settings where disease phenotypes are driven by complex cellular heterogeneity.

Beyond synthetic evaluations, we applied CTDP to real-world datasets spanning melanoma immunotherapy, COVID-19, and liver cirrhosis, thereby demonstrating its practical utility across diverse disease contexts. In melanoma, CTDP identified immune cell clusters enriched in responders, characterized by immune effector genes (e.g., PTPRC) and regulators (CREM, JUNB), highlighting potential biomarkers of immunotherapy efficacy. In COVID-19, CTDP uncovered multiple immune and epithelial populations associated with critical illness, marked by inflammatory gene expression and pathways such as neutrophil degranulation and interleukin signaling, with transcription factors like SPI1 and JUND implicated in immune dysregulation. In liver cirrhosis, CTDP pinpointed non-parenchymal cell types linked to fibrotic tissues and enriched in pathways related to inflammation and fibrogenesis, including platelet activation and IGF-binding protein regulation. Together, these results underscore CTDP’s ability to reveal phenotype-associated cellular subpopulations and their underlying molecular programs in complex disease settings.

Collectively, our results illustrate the ability of CTDP to not only identify phenotype-relevant cellular subsets with high accuracy but also to uncover the underlying molecular and regulatory features driving disease processes. Compared with previous approaches, CTDP offers several key advantages: (1) it directly models phenotype association via penalized regression, enabling explicit variable selection; (2) it incorporates rigorous statistical testing and FDR correction, enhancing result interpretability; and (3) it is applicable to diverse disease settings, from immune-mediated disorders to chronic fibrotic diseases.

Nevertheless, several limitations warrant further investigation. First, CTDP currently operates on discrete cell clusters and does not model continuous cellular trajectories or transitional states. Integration with pseudotime or RNA velocity analysis may extend its applicability to developmental or progressive disease models. Second, the reliance on predefined clustering may influence sensitivity to resolution and clustering algorithms. Future efforts to adapt CTDP to operate directly on the cell-level manifold (e.g., via graph-based smoothing or diffusion maps) could improve resolution. Lastly, although CTDP supports binary phenotypes, extending it to multiclass or continuous phenotypes (e.g., survival time, disease stage) would further expand its utility in clinical studies.

In conclusion, CTDP is a powerful and generalizable tool for dissecting the cellular architecture of complex diseases. By enabling robust identification of disease phenotype-associated cell populations and their regulatory programs, CTDP provides mechanistic insights and facilitates the development of cell-type-targeted therapeutic strategies.

## Materials and Methods

### scRNA-seq data and data preprocessing

We obtained the transcriptomic profiles of 16,291 immune cells from tumor samples of 48 melanoma patients treated with immune checkpoint inhibitors from the Gene Expression Omnibus (GEO) database (accession number GSE120575)^40^. In addition, single-cell transcriptomic data were collected from a published study, including nasopharyngeal, protected specimen brush, and bronchoalveolar lavage samples from 19 COVID-19 patients (8 moderate and 11 severe cases, as classified by the World Health Organization)^41^. To further expand the dataset, we also incorporated data from non-parenchymal cells in healthy and cirrhotic human livers reported in a previous study^42^. For each dataset, we utilized the raw count matrix and performed downsampling by cell type to balance the total number of cells between the two phenotypic groups.

The preprocessing of scRNA-seq data began with the selection of valid cells, retaining those with expression in at least three cells and containing a minimum of 200 detected genes. To remove low-quality cells, we calculated the proportion of mitochondrial gene expression for each cell and discarded cells with over 20% mitochondrial content. Gene expression data were normalized using the LogNormalize method, where the expression of each gene in a cell was divided by the total expression of that cell and multiplied by a scale factor of 10,000, followed by log-transformation. After normalization, we standardized gene expression such that each gene had a mean of 0 and a standard deviation of 1 across cells.

We then identified the top 2,000 highly variable genes across the dataset, which are considered to carry the most informative biological signals. Principal component analysis was performed based on these HVGs to extract major sources of variation. Cell-to-cell similarity was calculated using a principal component–based k-nearest neighbor graph constructed from the top 10 principal components. Based on this similarity graph, we applied graph-based clustering using the Louvain algorithm to identify distinct cellular subpopulations. Finally, dimensionality reduction and visualization were performed using Uniform Manifold Approximation and Projection (UMAP), enabling an intuitive representation of the relationships among identified cell subpopulations.

### Construction of simulated data

To evaluate the performance of disease-associated cell identification methods under controlled conditions, we constructed a simulated dataset based on a real single-cell RNA sequencing dataset consisting of 4,669 peripheral blood immune cells from two human donors^10^. After applying the same preprocessing pipeline described above, we selected one or more cell clusters and designated them as disease-associated target clusters, while the remaining clusters were treated as background cells.

Based on the distribution of cells in these real clusters, we assigned simulated sample labels to all cells. To regulate the proportion of cells with phenotype labels within the target clusters, we introduced a parameter called the mixing rate (α). For each cell in the target cluster, we assigned a sample label of 1 (indicating the presence of a phenotype under the simulated condition) with probability 1−*α*, and a label of 0 (background) with probability *α*. In this study, we set α to 0.1, meaning that approximately 90% of the cells in the target cluster were labeled as phenotype-positive, while the remaining 10% were treated as background. For cells in non-target clusters, sample labels were assigned randomly.

This labeling strategy enabled us to generate synthetic phenotype annotations for all cells, which were then used for downstream analyses. To ensure robustness and assess the consistency of each method, we repeated the experiment ten times, each time selecting different clusters as the target phenotype-associated clusters. The performance of each method was evaluated using the F1 score, which quantifies the agreement between predicted phenotype-associated cells and the ground truth labels.

### Description of CTDP

To systematically identify cell clusters associated with specific disease phenotypes, we developed a computational framework named CTDP (Identification of <u>C</u>ell <u>T</u>ypes associated with <u>D</u>isease <u>P</u>henotypes). CTDP integrates Lasso logistic regression with permutation-based significance testing, enabling robust feature selection in high-dimensional data while controlling for false discovery rates. The workflow of CTDP consists of three main steps: data encoding, model fitting, and statistical inference.

### Data encoding

The input data to CTDP includes two essential variables: (i) cluster, representing the cell cluster assignment of each cell, and (ii) phenotype, representing the sample-level disease label associated with each cell. To convert the categorical cluster labels into a format suitable for regression modeling, we apply one-hot encoding to the cluster variable. If the dataset contains *k* unique clusters {*C*_1_, *C*_2_, …, *C*_*k*_}, we define a binary design matrix *X* ∈ ℝ^*n*×*k*^, where *n* is the total number of cells. Each entry *X*_*ij*_ is defined as:

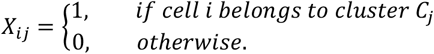

The phenotype variable is encoded as a binary response vector *Y* ∈ {0,1}^*n*^, where *Y*_*i*_ = 1 indicates that cell *i* originates from a sample with the phenotype of interest, and *Y*_*i*_ = 0 otherwise.

### Lasso logistic regression model fitting

To evaluate the association between each cluster and the disease phenotype, we fit a Lasso-regularized logistic regression model, which estimates the probability that a given cell belongs to the phenotype of interest using a linear combination of cluster indicators. The probability *p*_*i*_ that cell *i* is associated with the target phenotype is modeled as:

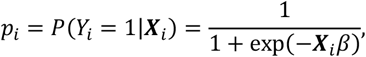

where ***X***_*i*_ ∈ ℝ^*k*^ is the one-hot encoded feature vector for cell *i*, and *β* ∈ ℝ^*k*^ is the coefficient vector representing the contribution of each cluster.

To enforce sparsity in the model and select only the most relevant clusters, we minimize the Lasso-penalized negative log-likelihood function:

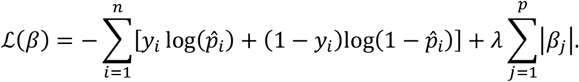

Here, *λ* ≥ 0 is the regularization parameter that controls the trade-off between model fit and sparsity. A larger *λ* leads to more coefficients being shrunk to zero. The optimal *λ*^∗^ is selected using cross-validation by minimizing the cross-validated classification error:

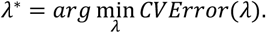

Once the model is fitted using *λ*^∗^, the non-zero coefficients *β*_*j*_ indicate clusters *C*_*j*_ that are most predictive of the phenotype.

### Permutation-based significance testing

To assess the statistical significance of each selected cluster, we perform a permutation test. The null hypothesis for each cluster *C*_*j*_ is that its coefficient *β*_*j*_ arises by chance. To construct a null distribution, we randomly permute the phenotype labels *Y* to obtain *Y*^*(b)*^ for each permutation *b* = 1,2, …, *B*, and refit the Lasso logistic regression model to obtain permuted coefficients 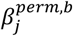. This results in a null distribution:

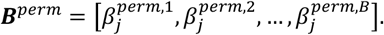

The empirical p-value *p*_*j*_ for each cluster *C*_*j*_ is then calculated as:

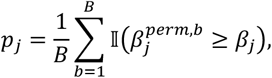

where 𝕀(·) is the indicator function. This p-value estimates the probability that a coefficient as large as *β*_*j*_ would be observed under the null.

To correct for multiple hypothesis testing, we apply the Benjamini– Hochberg procedure to the set of raw p-values {*p*_1_, *p*_2_, …, *p*_*k*_}. The false discovery rate (FDR) for each cluster is computed as:

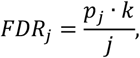

where *j* is the rank of *p*_*j*_ among all p-values sorted in ascending order. Clusters with adjusted FDR below a predefined threshold (e.g., 0.05) are considered significantly associated with the disease phenotype.

### Comparative methods for identifying phenotype-associated cell types

To validate the effectiveness and advantages of the proposed CTDP method, we compared its performance with several commonly used methods for identifying disease phenotype-associated cell types. The following methods were included in our comparative analysis:

DA-seq is a multiscale method based on a K-nearest neighbors (KNN) graph, designed to detect cell subpopulations with differential abundance between two biological conditions^43^. It computes a differential abundance score vector for each cell by examining its k-nearest neighbors across a range of k values. Based on the recommended settings in the DA-seq tutorial, the range of k was set from 50 to 500 with a step size of 50.

scDist is a statistical tool that leverages linear mixed-effects models to compare cell types across conditions while accounting for individual-level heterogeneity and technical noise in single-cell transcriptomic data ^44^. The method constructs an interpretable distance metric and approximates intergroup differences in a low-dimensional embedding. To reduce computational burden, we set the number of principal components to 8 in our implementation of scDist.

PENCIL is a recently developed method that utilizes a rejection learning strategy to identify high-confidence phenotype-associated cell subpopulations^45^. It combines a prediction function with a rejection function, improving classification accuracy by filtering out low-confidence predictions. All analyses using PENCIL were performed with default parameter settings.

### Differential expression and pathway enrichment analysis

In this study, differential expression analysis was performed using the Wilcoxon rank-sum test, as implemented in the FindMarkers function of the Seurat package. To identify significantly differentially expressed genes (DEGs), we applied a minimum absolute log2 fold change (|log2FC|) threshold of 0.5 and a Bonferroni-adjusted P-value cutoff of 0.05.

The resulting DEGs were subsequently subjected to pathway enrichment analysis using the Reactome database. This analysis was conducted with the ReactomePA package in R, which applies a hypergeometric test to assess the significance of each pathway. This approach enabled the identification of potential biological processes and signaling pathways associated with the identified DEGs.

### Gene regulatory network analysis

Gene regulatory network analysis was conducted using the SCENIC framework to explore the transcriptional regulation within the subpopulations identified by the CTDP method^46^. The analysis began with the identification of co-expression modules between transcription factors and their potential target genes using single-cell transcriptomic data. These co-expression modules were then refined with the R package RcisTarget, which predicts direct TF– target relationships by evaluating the enrichment of transcription factor binding motifs within the regulatory regions of co-expressed genes, based on genome-wide motif ranking databases. Following this, the activity of each regulon— defined as a transcription factor and its direct targets—was quantified across individual cells using AUCell, providing cell-specific regulon activity scores^47^. To identify key regulators associated with each subpopulation, we further calculated the Regulon Specificity Score (RSS), which is an entropy-based measure of the regulon’s activity specificity across clusters. Regulons with high RSS values were considered critical transcriptional regulators within the corresponding subpopulations.

## Supporting information

Supplementary_Table_1 Differentially expressed genes between CTDP-identified cells associated with immunotherapy responders versus all other cells

Supplementary_Table_2 Reactome pathways enrichment of upregulated genes in CTDP-identified cells associated with immunotherapy responders

Supplementary_Table_3 The specificity scores of regulons in CTDP-identified cells associated with immunotherapy responders

Supplementary_Table_4 Differentially expressed genes between CTDP-identified cells associated with severe COVID-19 versus all other cells

Supplementary_Table_5 Reactome pathways enrichment of upregulated genes in CTDP-identified cells associated with severe COVID-19

Supplementary_Table_6 The specificity scores of regulons in CTDP-identified cells associated with severe COVID-19

Supplementary_Table_7 Differentially expressed genes between CTDP-identified cells associated with cirrhosis versus all other cells

Supplementary_Table_8 Reactome pathways enrichment of upregulated genes in CTDP-identified cells associated with cirrhosis

Supplementary_Table_9 The specificity scores of regulons in CTDP-identified cells associated with cirrosis

## Data availability

The single-cell RNA sequencing datasets analyzed in this study are all sourced from publicly accessible repositories. The dataset of peripheral blood immune cells from two human donors is available from Synapse at https://www.synapse.org/Synapse:syn21041850^10^. Data for melanoma can be obtained from the Gene Expression Omnibus under accession number GSE120575^40^. The COVID-19 dataset is available via Figshare at https://doi.org/10.6084/m9.figshare.12436517^41^. The liver cirrhosis dataset can be accessed through the Edinburgh DataShare platform at https://datashare.ed.ac.uk/handle/10283/3433^42^.

## Code availability

The source code for CTDP, along with scripts necessary to reproduce the results presented in this study, is publicly available at the following GitHub repository: https://github.com/Chonghui-Liu/CTDP_project.

## Funding

This research was supported by the following funding: the National Natural Science Foundation of China (62472321,62225109,62450112); the Project Supported by Zhejiang Provincial Natural Science Foundation of China (Grant No. LTGY23H070004); the Project Supported by Quzhou municipal Science and Technology Project Foundation, Zhejiang Province, China (Grant No. 2022K55).

## Acknowledgements

We gratefully acknowledge the computing support provided by Northeast Forestry University, which facilitated the implementation of the algorithms presented in this study. We also extend our appreciation to the researchers who made their single-cell RNA sequencing datasets publicly available, enabling the analyses conducted in this work.

## Author contributions

C.L., G.W. and L.Y. conceived the overall research framework. The algorithm was developed and applied by C.L. Interpretation of the results was a joint effort by C.L., M.Z., Z.W., C.Y., Y.Z., Z.J., Y.Y., Z.L., and T.Z. The study was supervised by G.W. and L.Y. The manuscript was prepared by C.L., G.W. and L.Y., with input and critical revisions provided by all co-authors. All authors have read and approved the final version of the manuscript.

## Competing interests

The authors declare no competing interests.

## Notes

### Competing Interest Statement

The authors have declared no competing interest.

### Summary of Updates

The first author's affiliation has been updated

